# Differential pulmonary immunopathologic response of domestic sheep (Ovis aries) and bighorn sheep (Ovis canadensis) to Mycoplasma ovipneumoniae infection: a retrospective study

**DOI:** 10.1101/703249

**Authors:** Paige C. Grossman, David A. Schneider, Donald P. Knowles, Margaret A. Highland

## Abstract

*Mycoplasma ovipneumoniae* is a respiratory pathogen that can impact domestic sheep (*Ovis aries*; DS) and bighorn sheep (*Ovis canadensis*; BHS). Experimental and field data have indicated BHS are more susceptible than DS to developing polymicrobial pneumonia associated with *Mycoplasma ovipneumoniae* infection. We hypothesized that DS and BHS have a differential immunopathologic pulmonary response to *M. ovipneumoniae* infection. A retrospective study was performed using formalin-fixed, paraffin-embedded (FFPE) lung tissue from DS and BHS without and with *M. ovipneumoniae* detected in the lung tissue (n=8 per group). While each *M. ovipneumoniae* positive lung sample had microscopic changes typical of infection, including hyperplasia of intrapulmonary bronchus-associated lymphoid tissue (BALT) and respiratory epithelium, DS exhibited a more robust and well-organized BALT formation as compared to BHS. Immunohistochemistry was performed with antibodies reactive in FFPE tissues and specific for leukocyte and cytokine markers: T cell marker CD3, B cell markers CD20 and CD79a, macrophage markers CD163 and Iba1, and cytokine IL-17. Digital analysis was used to quantitate chromogen deposition in regions of interest (ROIs), including alveolar and bronchiolar areas, and bronchiolar subregions (epithelium and BALT). Main effects and interaction of species and infection status were analyzed by beta regression and Bonferroni corrections were performed on pairwise comparisons (*P_Bon_*<0.05 significance). Significant species differences were identified for bronchiolar CD3 (*P_Bon_*=0.0023) and CD163 (*P_Bon_*=0.0224), alveolar CD163 (*P_Bon_*=0.0057), and for IL-17 in each of the ROIs (alveolar: *P_Bon_*=0.0009; BALT: *P_Bon_*=0.0083; epithelium: *P_Bon_*=0.0007). Infected BHS had a higher abundance of bronchiolar CD3 (*P_Bon_*=0.0005) and CD163 (*P_Bon_*=0.0162), and alveolar CD163 (*P_Bon_*=0.0073). While IL-17 significantly increased with infection in BHS BALT (*P_Bon_*=0.0179) and alveolar (0.0006) ROIs, abundance in DS showed an insignificant decrease in these ROIs and a significant decrease in epithelial abundance (*P_Bon_*=0.0019). These findings support the hypothesis that DS and BHS have a differential immunopathologic response to *M. ovipneumoniae* infection.

## Introduction

*Mycoplasma ovipneumoniae* is a recognized agent associated with respiratory disease in members of the subfamilies Caprinae (sheep, goats, muskox) and Capreolinae (deer family members) [1–4]. Although clinically healthy domestic sheep (*Ovis aries*; DS) and bighorn sheep (*Ovis canadensis*; BHS) of both species can carry *M. ovipneumoniae*, anecdotal field reports and captive interspecies commingling and infection experiments provide evidence that BHS are more susceptible to *M. ovipneumoniae* associated pneumonia than are DS [5–7]. *M. ovipneumoniae* infection in DS primarily affects lambs, causing chronic respiratory disease, and a few reports describe infection in association with decreased growth [8, 9]. *M. ovipneumoniae* has been implicated as one of the bacterial pathogens associated with the complex and population-limiting phenomenon of epizootic pneumonia in BHS [5, 10, 11]. In order to mitigate interspecies transmission of respiratory pathogens, current policy decisions have opted for absolute separation of these two ovine species. Increasing restrictions on DS grazing on public land allotments and social pressures placed on private landowners has resulted in economic hardship and social upset. In order to formulate alternative mitigation strategies, mechanisms underlying the interspecies susceptibility differences to respiratory pathogens must be understood. Filling the current knowledge gap of the immunopathology associated with *M. ovipneumoniae* infections in DS and BHS will thus not only benefit animal health, but is also of socioeconomic and ecologic importance.

*M. ovipneumoniae* can serve as a primary pathogen, increasing the host’s susceptibility to other bacteria by adhering to respiratory epithelium and impairing cilia function which is necessary for mucociliary clearance [12–15]. Additionally, *M. ovipneumoniae* is also reported to adhere to the surface of macrophages, impairing phagocytosis of *M. ovipneumoniae* and potentially other bacteria that may be present [16, 17]. Altered immune functions such as these can increase host susceptibility to secondary pulmonary infections by other opportunistic pathogens residing in the upper respiratory tract. Such opportunistic pathogens, often reported in *M. ovipneumonaie* associated polymicrobial pneumonia, include: *Mannheimia haemolytica, Bibersteinia trehalosi, Pasteurella multocida, Trueperella pyogenes*, and *Fusobacterium* necrophorum [1, 18].

Histopathologic findings of *M. ovipneumoniae* infection in both DS and BHS include hyperplasia of intrapulmonary bronchus-associated lymphoid tissue (BALT) and respiratory epithelium, and mononuclear cell infiltrates surrounding (“cuffing”) bronchioles and within alveolar septa [2]. The immune response, including the cellular composition of the BALT hyperplasia and inflammatory infiltrates, to *M. ovipneumoniae* infection has yet to be characterized for DS and BHS. In other respiratory-associated *Mycoplasma* spp. infections, such as *Mycoplasma pulmonis* in mice and *Mycoplasma pneumoniae* in humans, severity of disease is reportedly dependent on the type of T helper immune response mounted by the host. For example, in both humans and mice, a Th1 (cell-mediated) response resulted in better management of infection, whereas a Th2 (antibody) response resulted in heightened pathology [19, 20]. Additionally, inhibition of the Th17 response in an IL-17 receptor knock-out mouse model resulted in higher bacterial load as compared to wild-type mice infected with *M. pulmonis* [20]. This led to the hypothesis that the reported interspecies susceptibility difference to *M. ovipneumoniae* associated polymicrobial pneumonia reported in DS and BHS is associated with a differential immunopathologic pulmonary response to *M. ovipneumoniae* infection. Therefore, the objective of this retrospective study was to qualitatively characterize and quantitatively compare the pulmonary immune responses of DS and BHS naturally infected with *M. ovipneumoniae*.

## Materials and Methods

### Lung tissue specimens

DS lung tissue was collected at University of Idaho Vandal Meats (Moscow, ID, USA) from lambs (estimated ages 7-12 months old) brought to slaughter between October 2016 and January 2018. DS lungs were grossly evaluated, the right cranial lobe removed, sterile swab samples collected from secondary bronchi, and a representative tissue sample from each animal was fixed in 10% neutral buffered formalin for 24 hours and a second sample was frozen fresh at −20°C.

BHS lung tissue sections (H&E and unstained sections for immunohistochemistry (IHC)) used in this study were attained from the Washington Animal Disease Diagnostic Laboratory (Pullman, WA, USA; WADDL), with permission granted by the submitting wildlife agency. BHS lung tissue sections were from archived formalin-fixed paraffin-embedded (FFPE) tissues collected from specimens submitted between November 2012 and February 2015 to the WADDL for gross, histopathologic, and bacteriologic analyses from animals (adults and lambs) that were either hunter harvested or culled for management purposes.

This retrospective study was carried out using lung tissues that were made available by opportunistic collection from sheep that were not maintained for research purposes and that died by means unrelated to this study. Thus, ethical approval for the use of animals in this study was not required.

### Histologic and microbial assessment of lung tissue

Fixed DS tissues were paraffin embedded, and 5 μm thick sections of both DS and BHS FFPE lung tissue were H&E stained at the WADDL. Stained slides from DS and BHS lungs were evaluated by light microscopy and the specimen was included in the infected (positive, POS) group if changes consistent with *M. ovipneumoniae* infection (BALT and bronchiolar epithelial hyperplasia) were present (Fig 1B), and *M. ovipneumoniae* was detected in the lung. To determine the *M. ovipneumoniae* infection status of each DS, DNA was extracted from bronchial swabs, and PCR and sequencing were performed, as previously described [3]. The *M. ovipneumoniae* infection status of each BHS was determined by real-time PCR performed at the WADDL at the time of specimen submission. Specimens were excluded from the study if histopathologic evaluation identified bronchopneumonia, indicative of secondary bacterial infection (Fig 1C). Eight sheep of each species that met the POS group inclusion criteria were selected for study. An additional 8 sheep of each species were selected for the “not detected” (ND) group based on the absence of *M. ovipneumoniae* DNA detection and the absence of histologic changes of infection or other abnormalities (Fig 1A). Three of the BHS specimens were from lambs (approximately 5-9 month old), 2 of which were placed in the “infected” group, and 13 were from adults. Light microscopic analysis was performed on H&E stained slides by a pathologist (MAH) to qualitatively characterize histopathologic changes. Aerobic bacterial culture on fresh (BHS) or fresh frozen (DS) lung specimens was performed at the WADDL to further assess for bacterial co-infections in each of the 32 animals included in this study.

**Fig 1.**
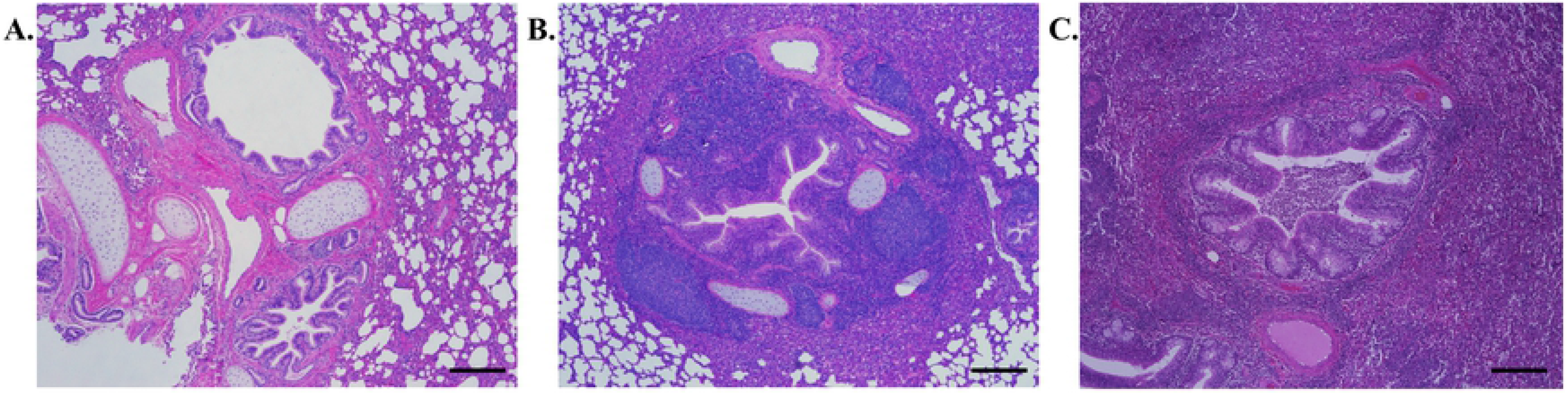
Histopathologic criteria for specimen selection. (A) Histologically normal lung selected for the *Mycoplasma ovipneumoniae* “not detected” (ND) group. (B) Lung section with histopathologic change consistent with *Mycoplasma ovipneumoniae* infection including bronchial associated lymphoid tissue (BALT) hyperplasia, epithelial hyperplasia, peribronchiolar lymphoid cuffing, and atelectasis, selected for infected group, selected for the *M. ovipneumoniae* positive (POS) group. (C) Lung section excluded from study showing histopathologic change consistent with bronchopneumonia (suspect secondary bacterial infection), including suppurative exudate in a bronchiole and alveolar inflammatory infiltrates. Representative domestic sheep lung tissue; H&E stain; scale bar = 250 μm.

### Immunohistochemistry

#### Protocol

FFPE tissue sections (3 μm) were placed on charged glass slides and baked at 56°C overnight before staining. Immunohistochemistry was carried out in a Ventana Discovery XT automated slide stainer using Cell Conditioner 1 (basic conditioner on medium setting) for antigen retrieval (eleven 4 minute incubations), DISCOVERY Antibody Block (4 minute incubation), DISCOVERY Universal Secondary Antibody cocktail (1 hour incubation), the DISCOVERY RedMap chromogen kit, and hematoxylin (4 minute incubation) and Bluing Reagent (4 minute incubation) counterstain (Roche, Ventana Medical Systems, Inc, Tucson, AZ, USA). Following antigen retrieval and preceding the antibody block step, Dako Dual Endogenous Enzyme Block was used to abolish endogenous enzyme activity (12 minute incubation; Agilent, Santa Clara, CA, USA). Each primary antibody was loaded onto the slide following the blocking step and incubated for 2 hours. After the staining was complete, slides were dipped 50 times in Dawn^®^ dishwashing liquid diluted in water, 10 times in tap water, 3 times in double distilled water, 30 times in 4 changes of 100% ethanol, and 50 times in 3 changes xylenes before cover-slipping.

#### Leukocyte and cytokine immunomarkers

Thirty-three antibodies were screened for use in this study (S1 Table). Antibodies that could be optimized (reactive in each species with specific staining and little to no background) included a T cell marker (CD3), B cell markers (CD20 and CD79a), macrophage markers (CD163 and Iba1), and one antibody for cytokine IL-17 (Table 1). Antibodies that could not be optimized were attempted at several different concentrations, with both basic (Cell Conditioner 1) and mildly acidic (Cell Conditioner 2) antigen retrieval methods (Roche, Ventana Medical Systems, Inc, Tucson, AZ, USA). While anti-CD45RO worked in DS tissue, it could not be optimized for use in the BHS tissues.

**Table 1.**
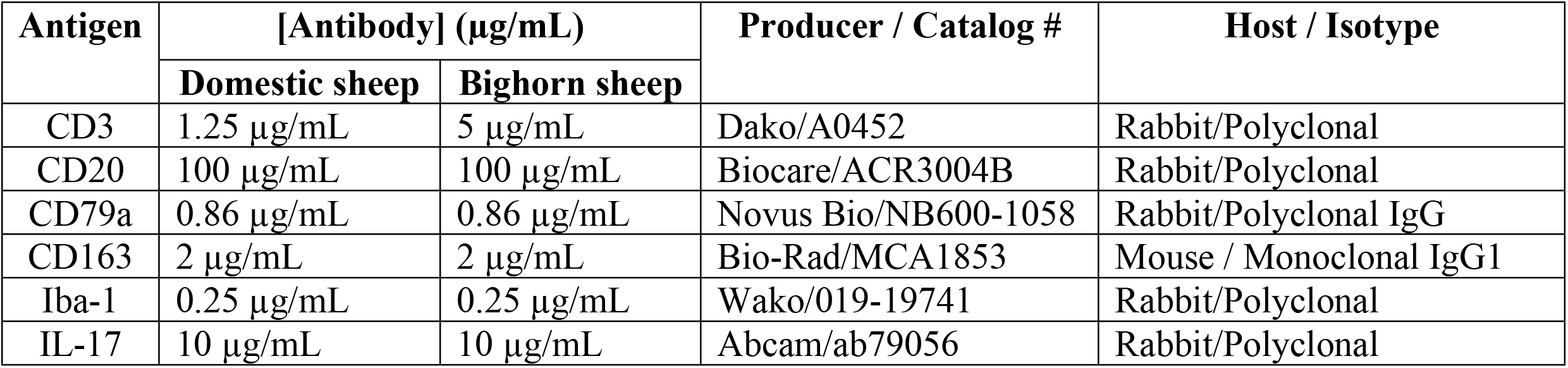
Antibodies optimized for immunohistochemically labeling formalin-fixed, paraffin embedded domestic and bighorn sheep tissue.

#### Control antibodies

Appropriate isotype negative control antibodies were used in each IHC run on representative BHS and DS tissue sections at the same concentration as the specific primary antibody. Non-immunized rabbit serum (X0903; Agilent, Santa Clara, CA, USA) was used as the negative control for the polyclonal antibodies and irrelevant epitope monoclonal IgG1, IgG2a, and IgG2b were used as the monoclonal negative controls (ab81032, ab18414, ab18457; Abcam, Eugene, OR, USA). Prior to use, total IgG was purified from the polyclonal negative control rabbit serum using Nab™ Protein A Plus Spin Kit, followed by dialysis against PBS containing 0.02% sodium azide using a Slide-A-Lyzer™ 3.5K MWCO Dialysis Cassette, following the manufacturer’s instructions (Thermo Scientific, Waltham, MA, USA). Protein concentrations of the non-immunized rabbit serum (polyclonal control) and the purified total IgG (polyclonal IgG control) were determined using the Pierce™ BCA protein assay kit, following the manufacturer’s instructions (Thermo Scientific, Waltham, MA, USA).

### Analysis

#### Tissue imaging

Immunolabeled slides were sent to the University of Washington Histology Imaging Core (Seattle, WA, USA) and scanned with a Hamamatsu NanoZoomer S360 (Hamamatsu Photonics, Bridgewater NJ, USA). Images were imported into Visiopharm software (Visiopharm, Hoersholm, Denmark) for quantitative analysis using previously described Visiopharm Image Analysis module settings [21]. Regions of interest (ROIs) were manually selected on each slide (S1 Fig). For five of the immunomarker antibodies (anti-CD3, anti-CD20, anti-CD79a, anti-CD163, and anti-Iba-1), the ROI included bronchi/bronchioles and the surrounding tissue including associated immune tissue (BALT, immune cell “cuffing”), but excluded bronchial cartilage and surrounding alveoli; these regions are referred to simply as “bronchiolar” ROIs (Image A in S1 Fig). Since IL-17 is produced by both immune cells and respiratory epithelial cells [22], the bronchiolar ROIs were each subdivided into 2 ROIs for quantifying IL-17 immunolabeling: bronchiolar epithelium and bronchiolar ROI’s excluding the epithelium, referred to as “epithelium” and “BALT”, respectively. Macrophage immunomarkers (CD163 and Iba1) and the IL-17 cytokine marker were also evaluated in alveoli and alveolar walls, this “alveolar” ROI was identified by selecting the whole lung tissue section and excluding bronchiolar ROIs (Image B in S1 Fig).

#### Statistical analysis

Leukocyte marker and cytokine abundances were quantitatively assessed as the ratio of immunolabeled area (μm^2^) to the total tissue area (μm^2^) of a given ROI. Non-specific chromogen deposition for each of the isotype negative control slides was similarly assessed, the ratio values of which were deducted from each test slide to provide the final area (μm^2^) of immunospecific chromogen deposition. Background correction resulted in bronchiolar chromogen ratios slightly less than zero for anti-CD20 (7 of 8 ND BHS) and anti-CD79a (2 of 8 ND and 1 of 8 POS BHS). Given that visual inspection of these slides confirmed minimal or no chromogen deposition other than background, a small constant was added to rescale all samples within each of these immunomarker groups, such that the greatest negative ratio values were equal to 1×10^−7^. The main effects and interaction of species (DS, BHS) and infection status (ND, POS) on the ratio measure of immunospecific chromogen deposition (a proportional response bounded by zero and one) was analyzed by beta regression (PROC GLIMMIX, logit link function; SAS version 9.4, SAS Institute Inc., Cary, NC, USA). The main effects and interaction of species and infection status on the mean area of epithelium ROI and mean area of BALT ROI per airway (total area of epithelium ROI or BALT ROI divided by the total number of bronchi and bronchioles), was also analyzed by beta regression (PROC GLIMMIX, identity link function; SAS version 9.4, SAS Institute Inc., Cary, NC, USA). A significant interaction term was parsed for all pairwise comparisons using the method of Bonferroni (*P_Bon_*). Significance was attributed to effects with P-values < 0.05. “Outlier” data were identified as values meeting two criteria: (1) the value was identified as a far outlier (i.e., values > 3x the interquartile range; PROC BOXPLOT, SAS version 9.4, SAS Institute Inc., Cary, NC, USA); and (2) if any of the absolute values of the Studentized residuals associated with a far outlier were greater than 3. Outliers so identified were excluded from the final analyses. Of note, outlier exclusion improved the apparent fit of the beta regression model but did not change the significance of main, interaction, or post-hoc effects of interest in any analysis. Although excluded from analysis, data identified as outliers are depicted as “x” within box plot presentations.

## Results

Qualitative assessment of H&E stained lung tissue sections determined that the samples from *M. ovipneumoniae* POS DS had a more densely cellular and organized BALT hyperplasia with secondary follicle formation as compared to the samples from POS BHS. Similarly, the samples from the ND DS had more centers of BALT observed than ND BHS. Epithelial hyperplasia was observed in both POS BHS and POS DS. Overall, notable differences in these observations were not noted between lamb and adult BHS lung tissue sections. Figure 2 illustrates examples of the most prominent BALT identified in 4 animals from each of the study groups.

**Fig 2.**
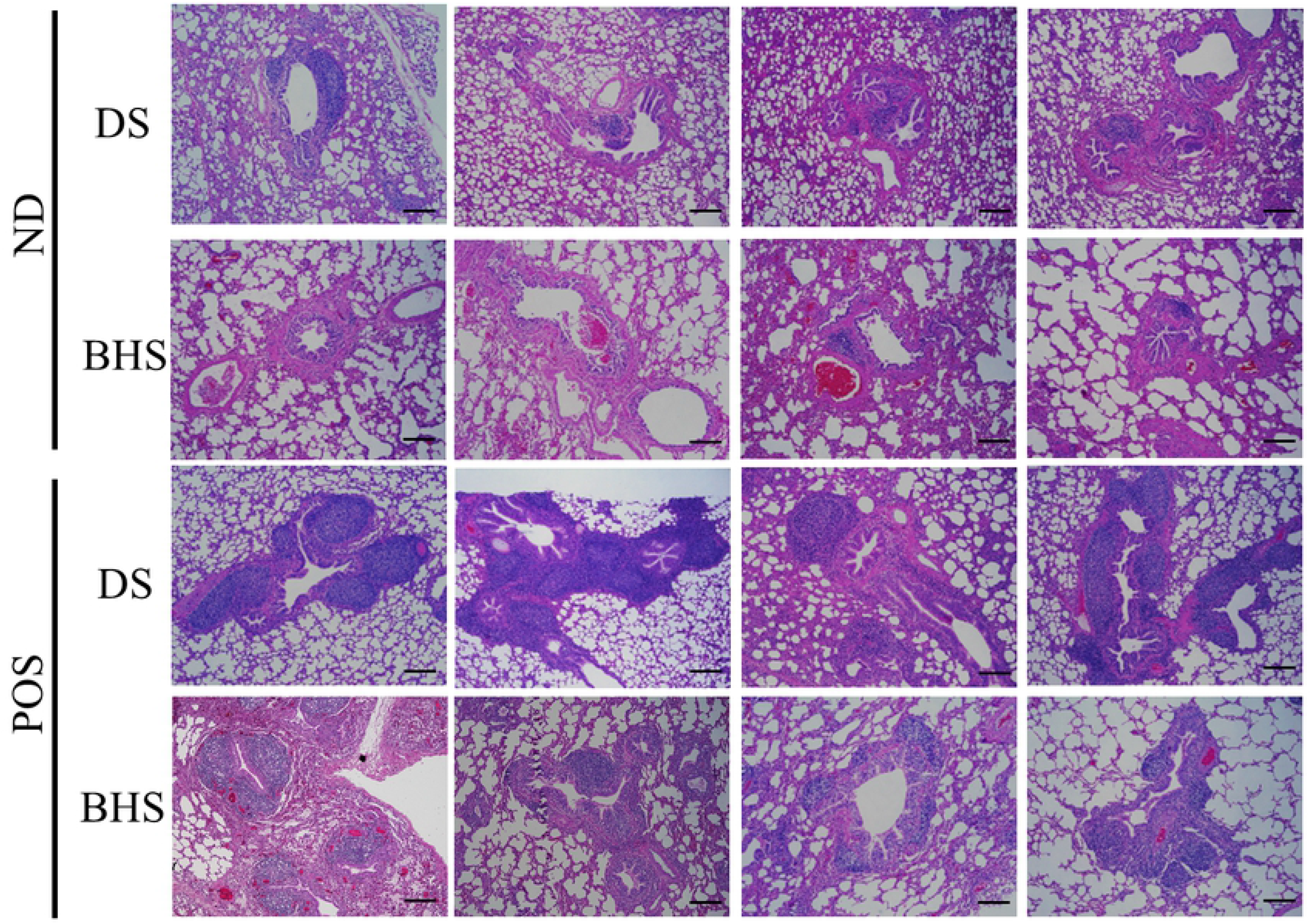
Representative images of lung tissue from domestic sheep and bighorn sheep without and with *Mycoplasma ovipneumoniae* infection. Light microscopic image examples of lung tissue from 8 domestic sheep (DS) and 8 bighorn sheep (BHS) in which *Mycoplasma ovipneumoniae* was not detected (ND) or detected (POS) in lung tissue. Images represent the most prominent areas of bronchus-associated lymphoid tissue (BALT) identified in animals within study. ND DS had more observed centers of BALT than ND BHS. Centers of BALT appeared densely cellular with secondary follicle formation in POS DS, as compared to the less organized and looser arrangement of the BALT observed in POS BHS samples. ND and POS BHS lambs are represented in column 1; all other BHS are adults. H&E stain; scale bar = 250 μm.

The results of digital quantification of BALT ROI and epithelial ROI areas are shown in Figure 3. Although there was no interaction between the effects of species and infection status, analysis supported the qualitative observation of BALT hyperplasia in POS animals, as infection was associated with a significantly larger overall mean area of BALT ROI per airway (*P_infection_*=0.0016). The estimate (mean and standard error) for the effect of infection status (POS vs ND) on the mean area (mm^2^) of BALT ROI per airway was 0.6216 ± 0.1769, representing a fold change of 1.862 (95% confidence limits = 1.295-2.677). The mean area of BALT ROI per airway was larger in BHS as compared to DS (*P_species_*=0.0079). The estimate for the effect of species (BHS vs DS) was 0.5064 ± 0.1766, representing a fold change of 1.659 mm^2^ (95% confidence limits = 1.155-2.384). The qualitative observation of epithelial hyperplasia in POS samples was not detected by digital quantification and analysis, as the area of epithelial ROI per airway was not significantly associated with effect of infection status (*P_infection_*=0.9244), nor was it associated with either the effect of species (*P_species_*=0.2306) or the interaction of these main effects (*P_interaction_*=0.0731).

**Fig 3.**
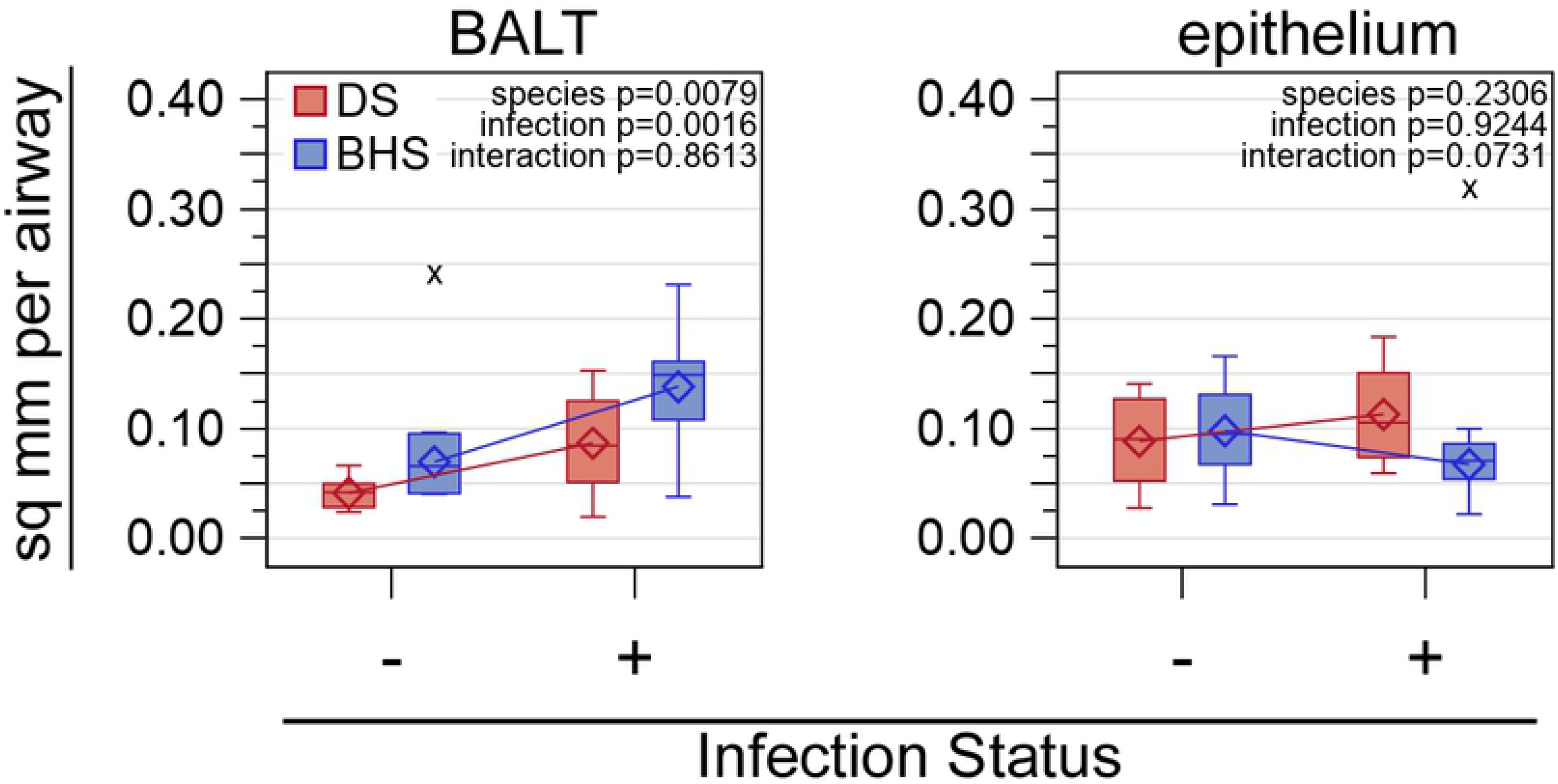
Quantitative analysis of bronchus-associated lymphoid tissue and bronchial/bronchiolar epithelium within lung tissue from domestic sheep and bighorn sheep without and with *Mycoplasma ovipneumoniae* infection. The Y-axes represent the area (mm^2^) of bronchial-associated lymphoid tissue (BALT) and bronchiole/bronchiolar epithelium, as indicated above each box plot. Infection status is represented on the X-axes by (-) and (+), indicating *Mycoplasma ovipneumoniae* was not detected or detected, respectively, in the domestic sheep (DS) and bighorn sheep (BHS). Box plot construction: box, interquartile range (IQR); open diamond, mean; horizontal line, median; vertical whiskers, data extending up to 1.5x IQR; open circle, data > 1.5x IQR; x, data identified as extreme values and excluded from formal analyses. Inset text: P-values for the main effects (species and infection status) and interaction term. The two outliers (“x”) are adults.

Analysis of the proportion of the area of chromogen deposition to the area of the ROI revealed significant interactions between the effects of species and infection status in the bronchiolar ROI for CD3 (*P*=0.0023) and CD163 (*P*=0.0224), and in the alveolar ROI for CD163 (*P*=0.0057). Infection was associated with significantly higher bronchiolar chromogen deposition for CD3 in BHS (~7.1-fold; *P_Bon_*<0.0001) but a significant difference was not detected between ND and POS DS (*P_Bon_*=0.1989). Similarly, infection was associated with significantly higher bronchiolar CD163 chromogen deposition in BHS (~4.0-fold; *P_Bon_*=0.0064), but a significant difference was not detected between ND and POS DS (*P_Bon_*=1.0000). Although the interaction term for the proportion of bronchiolar CD20 chromogen deposition was not considered significant (*P_interaction_*<0.0527), the overall proportion was significantly less in BHS than in DS (*P_species_*<0.0001) and significantly higher in association with infection in both species (*P_infection_*<0.0001). The proportion of bronchiolar CD79a chromogen deposition was higher with infection (*P_infection_*=0.0127) but significant effects of species were not detected (*P_species_*=0.1827; *P_interaction_*=0.6063). Bronchiolar Iba1 chromogen deposition was not associated with significant main effects of species (*P_species_*=0.0871) or infection status (*P_infection_*=0.5642), nor with significant interaction of these terms (*P_interaction_*=0.4691).

Within the alveolar ROI, a significant interaction of species and infection status was detected for CD163 (*P_interaction_*=0.0057). Infection was associated with higher alveolar chromogen deposition for CD163 in BHS (~4.3-fold; *P_Bon_*=0.0062) but no difference was detected between ND and POS DS (*P_Bon_*=1.0000). Alveolar Iba1 chromogen deposition was not associated with significant main effects of species (*P_species_*=0.2336) or infection status (*P_infection_*=0.9897), nor with significant interaction of these terms (*P_interaction_*=0.9426).

For IL-17, significant interactions of species and infection status were detected in each of three ROI (BALT, *P_interaction_*=0.0083; bronchiolar epithelium, *P_interaction_*=0.0007; alveolar, *P_interaction_*=0.0009). Within the BALT ROI, IL-17 chromogen deposition was significantly higher in association with infection in BHS (~2.2-fold, *P_Bon_*=0.0179) but no difference was detected between ND and POS DS (*P_Bon_*=1.0000). Similarly, alveolar IL-17 chromogen deposition immunolabeling was significantly higher in association with infection in BHS (~4.3-fold, *P_Bon_*=0.0006) but no significant difference was detected between ND and POS DS (*P_Bon_*=1.0000). In contrast, infection was associated with significantly lower IL-17 chromogen deposition within bronchiolar epithelium in DS (~2.6 fold, *P_Bon_*=0.0019), but no difference was detected between ND and POS BHS (*P_Bon_*=1.0000).

Representative images of immunolabeled and negative isotype control sections are shown in Figure 4. The results of digital quantification for all immune cell markers within the bronchiolar ROI (Fig 5A) and for macrophage markers within the alveolar ROI (Fig 5B) are summarized as box-and-whiskers plots. Quantification for the cytokine, IL-17, immunomarker specifically within the BALT, epithelium, and alveolar ROIs is also summarized as box-and-whiskers plots (Fig 6). Outliers, depicted by “x” in Figures 4 and 5, identified for the BHS are adult animals, except for the POS BHS CD20 outlier which is a lamb. The intraspecies and interspecies fold changes (odds ratios) in immunospecific chromogen deposition associated with infection status are provided in Table 2.

**Fig 4.**
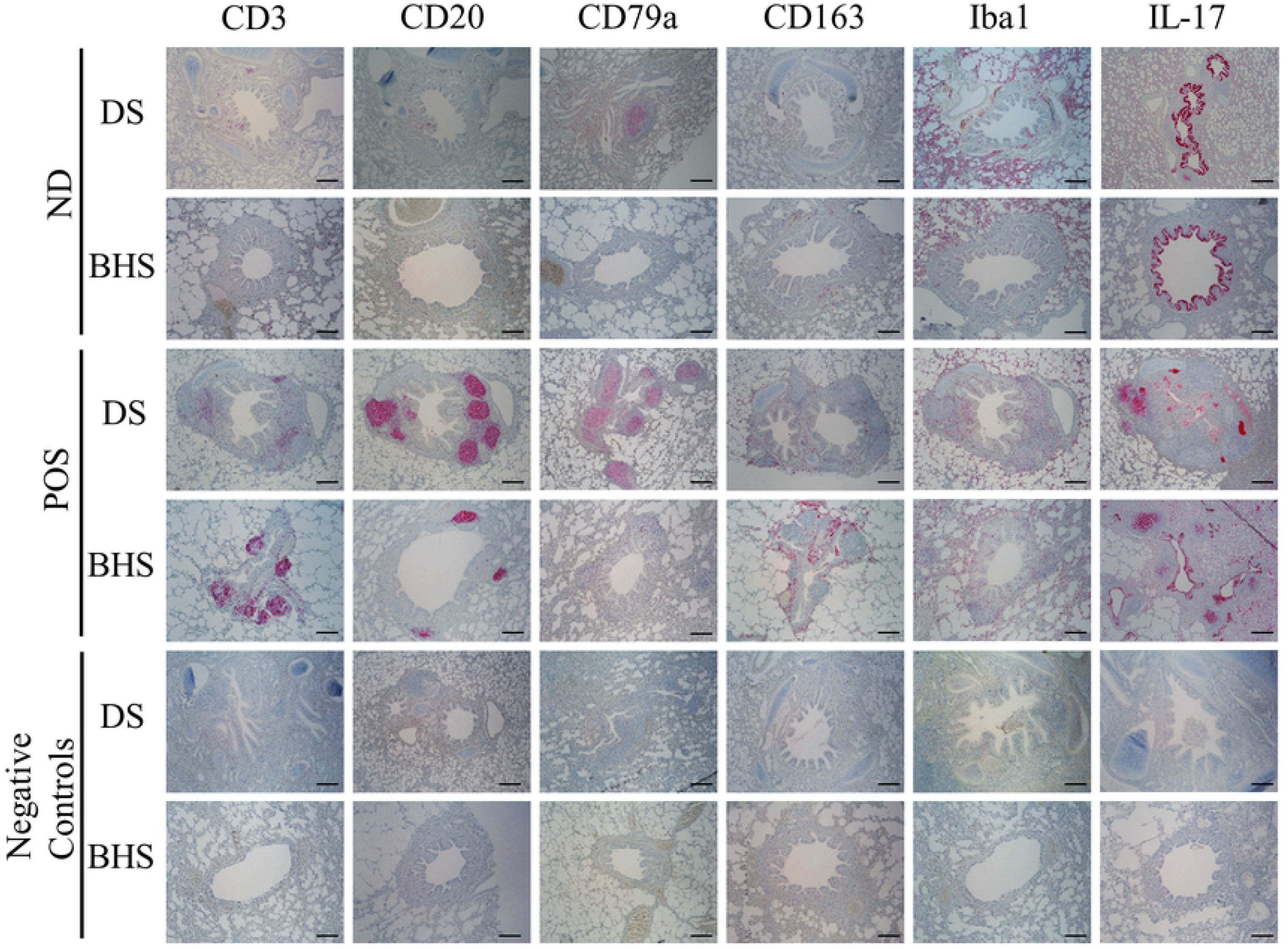
Representative immunolabeled lung tissue sections from domestic sheep and bighorn sheep without and with *Mycoplasma ovipneumoniae* infection. Light microscopic images of immunolabeled (top 4 rows) and negative control (bottom 2 rows) lung tissue, including bronchi/bronchioles and surrounding tissue, from domestic sheep (DS; rows 1, 3, 5) and bighorn sheep (BHS; rows 2, 4, 6) that had *Mycoplasma ovipneumoniae* either not detected (ND; top 2 rows) or detected (POS; middle 2 rows) in lung tissue. Red chromogen deposits represent detection of leukocyte markers and a cytokine, as indicated by column headings. Counter stain: hematoxylin and Bluing Reagent; scale bar = 250 μm.

**Fig 5.**
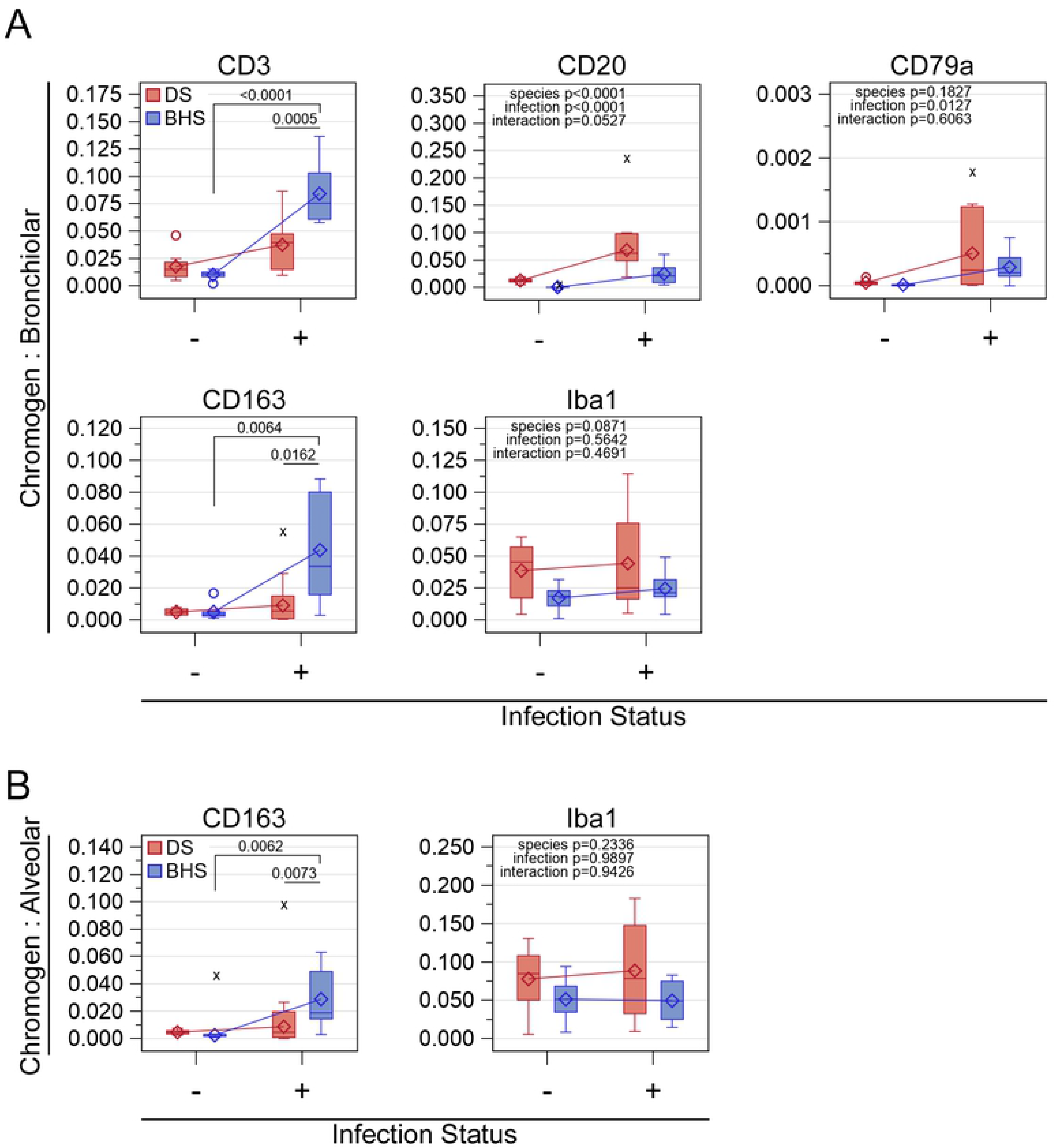
Quantitative analysis of leukocyte markers within lung tissue from domestic sheep and bighorn sheep without and with *Mycoplasma ovipneumoniae* infection. The Y-axes represent ratio of the area of immunospecific chromogen to total bronchial/bronchiolar tissue (“bronchiolar”) area (A), and to total alveolar area (B). Areas of chromogen deposition represent detection of leukocyte markers, as indicated above each box plot. Infection status is represented on the X-axes by (-) and (+), indicating *Mycoplasma ovipneumoniae* was not detected or detected, respectively, in the domestic sheep (DS) and bighorn sheep (BHS). Box plot construction: box, interquartile range (IQR); open diamond, mean; horizontal line, median; vertical whiskers, data extending up to 1.5x IQR; open circle, data > 1.5x IQR; x, data identified as extreme values and excluded from formal analyses. Statistical bars: significant difference between infection statuses within the indicated species (bar with drop lines) or between species within an infection status (bar without drop lines). Inset text: statistical significance of the main effects when the interaction term was insignificant. All data used for analysis are provided for review (S2 Table). The (-) BHS outliers include 1 lamb (CD20) and 1 adult (CD163).

**Fig 6.**
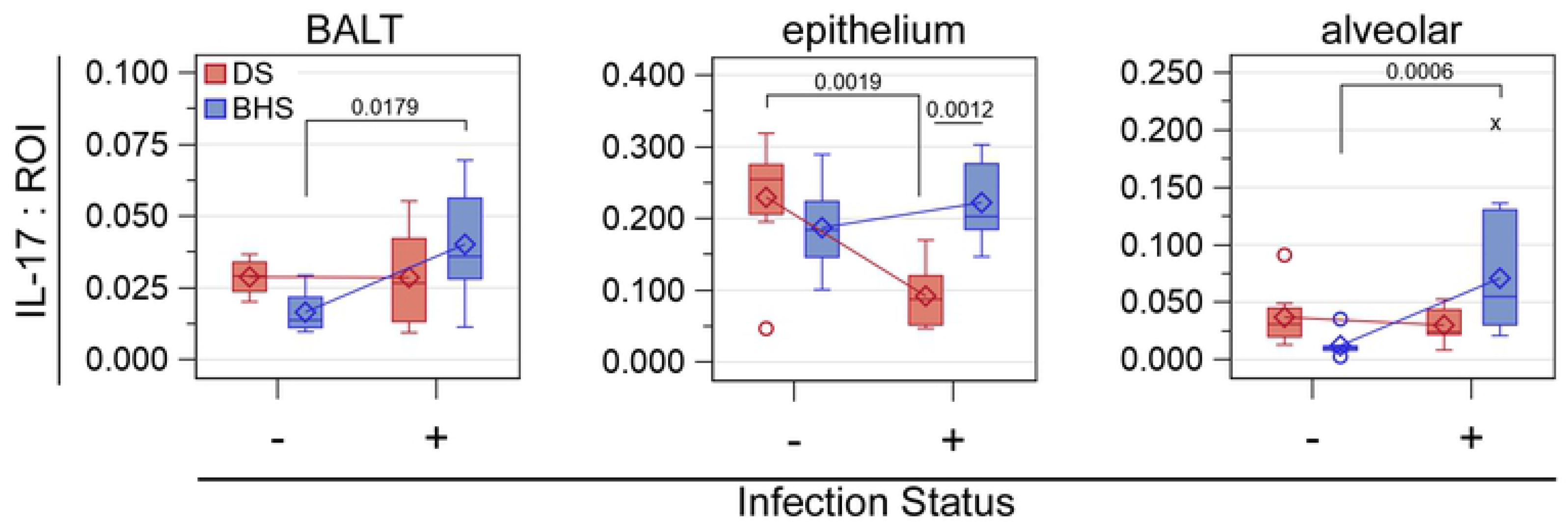
Quantitative analysis of cytokine IL-17 within lung tissue from domestic sheep and bighorn sheep without and with *Mycoplasma ovipneumoniae* infection. The Y-axes represent ratio of the area of IL-17 immunospecific chromogen deposition to the region of interest (ROI) as indicated above each boxplot. Infection status is represented on the X-axes by (-) and (+), indicating *Mycoplasma ovipneumoniae* was not detected or detected, respectively, in the domestic sheep (DS) and bighorn sheep (BHS). Box plot construction: box, interquartile range (IQR); open diamond, mean; horizontal line, median; vertical whiskers, data extending up to 1.5x IQR; open circle, data > 1.5x IQR; x, data identified as an extreme value and excluded from formal analyses. Statistical bars: significant difference between infection statuses within the indicated species (bar with drop lines) or between species within an infection status (bar without drop lines). Interaction is significant (*P_interaction_*<0.05) for each ROI. All data used for analysis are provided for review (S2 Table). The single BHS outlier is an adult.

**Table 2.** Intraspecies and interspecies fold change in chromogen deposition in domestic sheep and bighorn sheep without and with *Mycoplasma ovipneumoniae* detected in lung tissue.

| Antigen | Region of Interest | Intraspecies POS:ND<br>Fold Change (CL<sub>Bon</sub>)<br>*P*<sub>Bon</sub> |  | Interspecies BHS:DS<br>Fold Change (CL<sub>Bon</sub>)<br>*P*<sub>Bon</sub> |  |
| --- | --- | --- | --- | --- | --- |
|  |  | BHS | DS | ND | POS |
| CD3 | bronchiolar | 7.061 (3.165, 15.752)<br><0.0001 | 1.880 (0.845, 4.184)<br>0.1989 | 0.701 (0.272, 1.807)<br>1.0000 | 2.633 (1.450, 4.782)<br>0.0005 |
| CD20 | bronchiolar | 11.990 (3.378, 42.551)<br><0.0001 | 4.316 (1.969, 9.463)<br><0.0001 | 0.122 (0.034, 0.433)<br>0.0004 | 0.339 (0.162, 0.708)<br>0.0017 |
| CD79a | bronchiolar | 2.854 (0.755, 10.792)<br>0.1990 | 2.043 (0.563, 7.418)<br>0.7578 | 0.544 (0.141, 2.107)<br>1.0000 | 0.760 (0.222, 2.608)<br>1.0000 |
| CD163 | bronchiolar | 3.970 (1.361, 11.584)<br>0.0064 | 0.987 (0.287, 3.395)<br>1.0000 | 0.888 (0.266, 2.969)<br>1.0000 | 3.572 (1.192, 10.703)<br>0.0162 |
|  | alveolar | 4.325 (1.394, 13.420)<br>0.0062 | 0.728 (0.211, 2.507)<br>1.0000 | 0.705 (0.204, 2.438)<br>1.0000 | 4.186 (1.358, 12.900)<br>0.0073 |
| Iba1 | bronchiolar | 1.377 (0.486, 3.902)<br>1.0000 | 0.964 (0.390, 2.384)<br>1.0000 | 0.543 (0.199, 1.480)<br>0.5686 | 0.775 (0.299, 2.009)<br>1.0000 |
|  | alveolar | 0.980 (0.364, 2.638)<br>1.0000 | 1.014 (0.410, 2.511)<br>1.0000 | 0.762 (0.295, 1.971)<br>1.0000 | 0.737 (0.285, 1.907)<br>1.0000 |
| IL-17 | BALT | 2.153 (1.102, 4.206)<br>0.0179 | 0.858 (0.458, 1.609)<br>1.0000 | 0.576 (0.289, 1.151)<br>0.1895 | 1.447 (0.791, 2.647)<br>0.5615 |
|  | alveolar | 4.331 (1.733, 10.828)<br>0.0006 | 0.868 (0.384, 1.965)<br>1.0000 | 0.414 (0.160, 1.072)<br>0.0820 | 2.063 (0.959, 4.438)<br>0.0723 |
|  | epithelium | 1.251 (0.702, 2.231)<br>1.0000 | 0.378 (0.192, 0.741)<br>0.0019 | 0.832 (0.465, 1.487)<br>1.0000 | 2.756 (1.407, 5.397)<br>0.0012 |

The proportion of chromogen deposition within each region of interest relative to *Mycoplasma ovipneumoniae* infection status
(infected (POS) versus not detected (ND) for domestic sheep (DS) and bighorn sheep (BHS). Changes of interest included intraspecies
comparisons (ND BHS compared to POS BHS; ND DS compared to POS DS) and interspecies comparisons (ND BHS compared to ND DS; POS BHS compared to POS DS). Odds ratios are reported as fold change and Bonferroni adjusted 95% confidence limits are reported parenthetically; Bonferroni adjusted P-values ( $P_{Bon}$ ) <0.05 are considered significant.

## Discussion

The goal of this retrospective study was to test the hypothesis that there is a phenotypic interspecies difference in the pulmonary immune response exhibited by DS and BHS naturally infected with *M. ovipneumoniae*. Examining the response in naturally infected animals, living and managed as they otherwise exist, was a primary criterion of this project. This criterion was set based on the potential impact that captive environments can have on wild species, specifically alterations in immune responses secondary to stress or potential suboptimal nutrition. Additionally, this retrospective study design supports animal reduction in research, which supports the United States legislative mandate to incorporate the three Rs (reduction, refinement, and replacement) into research [23]. Since study design limited the available specimens for BHS to archived FFPE tissue, the opportunistically collected DS lung tissues were similarly fixed and processed. While formalin fixation followed by processing into paraffin blocks is an excellent method for maintaining tissue architecture for light microscopic examination, and is a common way to archive tissues, this processing can mask protein epitopes and thus limit antibodies for use in IHC. However, identifying antibodies for use in FFPE tissue is relevant and important, as already mentioned, many archived tissues are preserved in this manner and can serve as a valuable resource to researchers while reducing animal use in research.

Results of this study indicate that BHS respond to *M. ovipneumoniae* infection with a significantly more prominent T cell response as compared to DS. Multiple attempts were made to characterize the type(s) of T cells present in the bronchiolar ROIs (S1 Table) without success, likely due to the described limitations of using FFPE tissue. Regardless, the significantly higher chromogen deposition of T cell immunomarker CD3 in POS BHS, and B cell marker CD20 in POS DS (Table 2) substantiates the qualitative observation that POS BHS had dispersed or loosely arranged BALT and little to no follicle formation as compared to the more densely cellular, secondary follicle containing BALT observed in POS DS.

While the abundance of the macrophage marker CD163 increased significantly with infection in BHS bronchiolar and alveolar ROIs, as compared to DS which exhibited no change with infection in either ROI, the pan-macrophage marker, Iba-1, remained similar with infection in both species. CD163+ macrophages are activated along the alternative (M2) pathway by both pro-inflammatory Th2 cytokines and anti-inflammatory (glucocorticoids) stimuli [24]. Further investigation is required to determine which of these activation pathways predominates with infection in BHS; however, this result may be situation dependent and difficult to repeat experimentally if activation was by the anti-inflammatory pathway, secondary to unrecognized environmental stressors.

Perhaps the most interesting results from this study were for cytokine IL-17. IL-17 is expressed by Th17 cells, as well as NK cells, and neutrophils, and has been reported in other species to be expressed within pulmonary epithelium [22, 25]. IL-17 is a secreted cytokine that binds pulmonary epithelial cells inducing mucin production and stimulates neutrophil recruitment to the site of infection [26]. In this study, IL-17 significantly increased with infection in BHS BALT and alveolar ROIs while remaining similar in these ROI’s for DS. However, IL-17 significantly decreased with infection within the bronchial/bronchiolar epithelium in DS, while BHS had no detected change, remaining at an abundance similar to that of the uninfected DS. In murine studies, mice that were unable to produce an IL-17 response had depleted amounts of neutrophils and larger numbers of *M. pulmonis* present in the lung [20]. This suggests that IL-17 can contribute to an effective immune response to this pathogen through recruitment of neutrophils, although exuberant recruitment of inflammatory cells, particularly neutrophils, to pulmonary tissue can cause host cell damage that outweighs the benefit. The abundance of IL-17 in both uninfected DS and BHS and infected BHS pulmonary respiratory epithelium, in the absence of neutrophil influx, may indicate the IL-17 was produced locally but largely remained intracellular. IL-17 stimulated neutrophil recruitment may be of particular interest in BHS, as previous research supports higher neutrophil recruitment with pulmonary inflammation in BHS as compared to DS [27]. Additionally, BHS neutrophils have been shown to have heightened sensitivity to the cytotoxic effects of bacterial toxins, such as leukotoxin produce by *M. haemolytica* and *F. necrophorum* [18, 28, 29]. Given the importance of these leukotoxin producing bacteria in polymicrobial ovine pneumonia, exacerbation of neutrophil recruitment by IL-17 may in part explain the heightened morbidity and mortality described in BHS. In addition to IL-17 induced mucin production, experimental *M. pneumoniae* infection in mice has been shown to stimulate mucin production through toll-like receptor 2 [30]. Excess production of mucin in addition to mucociliary clearance impediment may act synergistically to enhance an environment favorable to colonization of bacteria that, under normal healthy lung conditions, are aspirated but then quickly cleared from the lungs.

Limitations and potential confounding factors to acknowledge in this study include the unknown time course of infection, pulmonary bacterial load in each animal, precise ages of the sheep, and precise site of sample collection for the BHS. Although the ages varied more in BHS, as compared to the DS, there was no indication that age impacted the study, as just 1 of the 5 outlier BHS data points was from a lamb, and no consistent lamb versus adult BHS trends were noted in the raw data (S2 and S3 Tables). Although the collection of lung tissue between species was considered an uncontrolled aspect of this retrospective study and a more bronchi were present in the DS samples, interspecies evaluation of the total number of airways, number of bronchioles, and total tissue area for the specimens used in this study were not significantly different (S2 Table). The larger number of bronchi in DS samples likely indicates specimens were collected closer to a primary bronchus than were the BHS tissues.

These data begin to define the immune responses found in the lungs of DS and BHS in the presence of *M. ovipneumoniae* infection. Especially interesting are the findings concerning comparative IL-17 levels. Critically, future work needs to address the interactions of environmental factors (*e.g*. stress, nutrition), host factors (*e.g*. genetics), and other potentially synergistic pathogens that may influence the immune response and induction of pneumonic disease associated with *M. ovipneumoniae* in DS and BHS.

## Acknowledgments

We wish to thank Dr. Charles Frevert, Brian Johnson, and Megan Larmore at the University of Washington Histology and Imaging Core Laboratory for their assistance; with special thanks to Megan for her excellent guidance in whole slide scanning and quantitative analysis. We are grateful to Dr. William Davis with the Washington State Monoclonal Antibody Laboratory for providing multiple monoclonal antibodies tested for use in this study. We thank the employees at the University of Idaho Vandal meets for providing the means for domestic sheep sample collection. We also thank Lori Fuller for immunohistochemistry guidance and Nicholas Durfee for general laboratory assistance.

---

**S1 Table. Antibodies screened for immunolabeling formalin-fixed, paraffin-embedded tissue from domestic sheep and bighorn sheep.**

**S2 Table. Measures of lung tissue thin section area by study subject.** Bighorn sheep lambs are highlight green. Outliers are bolded with an asterisk in the sheep ID column. Species: domestic sheep, DS; *Mycoplasam ovipneumoniae* infection status: not detected, ND; detected, POS.

**S3 Table. Measures of proportional chromogen deposition area within thin section by study subject.** Bighorn sheep lambs are highlight green. Outliers are bolded with an asterisk in the sheep ID column. Region of interest, ROI; species: domestic sheep, DS; bighorn sheep, BHS; *Mycoplasma ovipneumoniae* infection status: not detected, ND; detected, POS.

**S1 Fig. Tissue region of interest selection for analysis using Visiopharm software.** (A) Bronchiolar region selection (encircled by blue dashed line). (B) Alveolar regions selected by deselection of bronchiolar regions (encircled by black dash lines) from total lung tissue area (encircled blue dashed line); lymph node present on slide is excluded from selection. (C) Original scanned image zoomed in on alveolar spaces. (D) Red marks summed = positively stained area; green highlighted tissue = area counterstained. Ratio of immunolabeled area = Red ÷ (Red + Green).

**S2 Fig. CD79a immunohistochemical staining in bighorn sheep tissues.** Higher magnification of anti-CD79a antibody immunolabeled (A) bighorn sheep lymph node used for antibody optimization and validation, and (B) bronchus-associated lymphoid tissue from one of the bighorn sheep in this study.

